# Cell density couples tissue mechanics to control the elongation speed of the body axis

**DOI:** 10.1101/2023.12.31.573670

**Authors:** Changqing Lu, Joana M. N. Vidigueira, Christopher Chan Jin Jie, Alicja Maksymiuk, Fengzhu Xiong

## Abstract

The vertebrate body forms by addition of new tissues at the posterior end. This leads to body axis elongation which balances the anterior segmentation process to produce the stereotypic body plan. How elongation speed is constrained remains unknown. Here we utilised modeling and tissue force microscopy on chicken embryos to show that cell density of the posterior presomitic mesoderm (pPSM) dynamically modulates elongation speed in a negative feedback loop. Elongation alters the cell density in the pPSM, which in turn controls progenitor cell influx through the mechanical coupling of body axis tissues. This enables responsive cell dynamics in over- and under-elongated axes that consequently self-adjust speed to achieve long-term robustness in axial length. Our simulations and experiments further suggest that cell density and FGF activity act synergistically to drive elongation. Our work supports a simple mechanism of morphogenetic speed control where the cell density relates negatively to progress, and positively to force generation.

## INTRODUCTION

During body axis formation in vertebrate embryos, a progenitor zone at the posterior end of the axis progressively adds new cells to elongate tissues from different germ layers, such as the neural tube and the presomitic mesoderm. These tissues subsequently undergo morphogenesis and differentiation, such as segmentation, to create the initial body plan of the animal. In avian embryos, the posterior presomitic mesoderm (pPSM), which receives new cells from the tailbud, produces an expansive force that is a key driver of body axis elongation^1–3^. Elongation moves an FGF gradient posteriorly, which controls the boundary placement of new segments in conjunction with the oscillatory genes of the segmentation clock^4^. As the period of the segmentation clock is insensitive to body size^5,6^ or elongation progress^7,8^, the number and size consistency of segments depend on stable, robust elongation. However, how the elongation speed is constrained remains unknown.

The elongation speed is a collective phenotype of multiple processes such as new cell addition and tissue-force-driven deformation, and their interactions^9^. The cellular mechanisms leading to the expansive forces of the pPSM that drive elongation remain unclear, but have recently been proposed to rely on random cell motility and/or extracellular-matrix-driven swelling downstream of FGF signaling^1,3,10^, and in the longer term, proliferation^11^. All of these processes depend on the cell density of the pPSM, making it a potential direct regulator of the elongation speed. Indeed, the avian presomitic mesoderm (PSM) exhibits a characteristic anterior to posterior cell density gradient (from ∼10.2k/mm^2^ in the anterior PSM to ∼6.5k/mm^2^ in the pPSM^2^) during elongation. The pPSM’s particularly low cell density as compared to its surrounding tissues is associated with its mesenchymal cell state and tissue expansion^1^. The resulting lateral compressive stresses on the axial tissues promote their elongation^2,12^, which in turn promotes the lateral movement of midline PSM progenitors into the pPSM (the progenitor cell influx), completing an engine-like positive feedback loop^2^. Cell density is therefore regulated by both pPSM expansion and progenitor influx, which work in opposite directions, making cell density a natural node for negative feedback from the perspective of an integrated system (Figure 1A).

**Figure 1.**
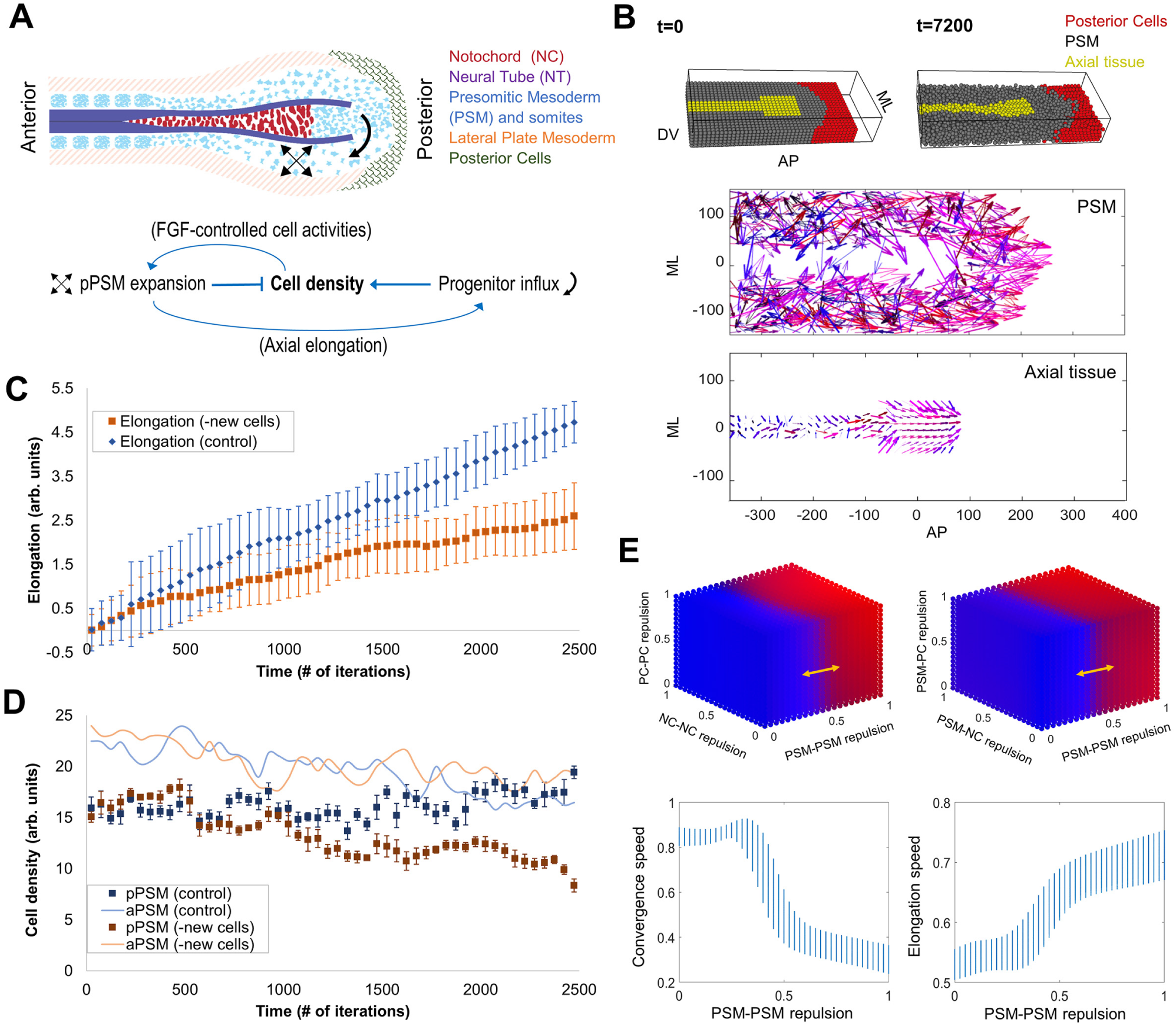
Modeling axis elongation and pPSM cell density dynamics. **A.** Schematic of the tissue organization of the posterior body axis. The network diagram depicts the interactions that alter cell density in the pPSM. **B.** Visualization of PhysiCell 3D tissue layout generated in PhysiCell Studio, t (time) indicates the simulation minutes. A portion of the PSM (grey) has been made invisible, allowing for the shape of the axial tissue (yellow) to be seen. DV, dorsal-ventral axis, dorsal to the top; AP, antero-posterior axis, anterior to the left; ML, medio-lateral axis, axial tissue is the medial. Bottom plots show the vectors of cell displacements in a representative simulation. Arrow length represents the displacement of cells (magnitude) and direction (also color-coded, blue represents convergence behaviours, red represents elongation behaviours, thickness indicates displacements in Z, i.e., DV). Both the axial tissues and pPSM are observed to converge along the ML axis and elongate along the AP axis towards the posterior. The anterior regions of the tissues are dominated by convergence (lateral to medial) movements (mostly blue arrows). The posterior region is dominated by elongation (anterior to posterior, mostly red arrows) and convergence extension (mostly purple arrows). **C.** Simulation of elongation in the 2D model without new cell addition from the progenitor domain. The total progress of elongation was measured over simulation time. 20 simulations were performed in each condition and 50 adjacent time points were binned to generate the plot. Error bars are ±SD of the binned mean. **D.** Cell density in the simulated PSM corresponding to panel **C**. Cells were counted in grids specified (anterior and posterior PSMs are labelled as aPSM and pPSM, respectively) in the simulated PSM as a function of distance to the axis end. A decrease in cell density in the pPSM of the simulations lacking new cell addition is observed. The density rise of the pPSM towards the end was a simulation artefact as the axial cells which do not divide in the model run out, causing pPSM cells to accumulate on the midline. **E.** Parameter space sweep of cell-cell interactions using a simplified 2D model (without new cell addition). Each cube compares 2 independent scaled parameters with PSM-PSM repulsion on their effect on tissue movement (blue, convergence dominated; red, elongation dominated), accounting for all 5 parameters used in the model. Bottom plots show the impact of PSM-PSM repulsion in comparison to NC-NC and PC-PC repulsions (shown as the vertical range of the blue bars). PSM-PSM repulsion appears to be the determinant factor of the behaviour of the system, with a sharp transition boundary from convergence dominated to extension/elongation dominated cell movements around 0.5. PC, posterior cells; NC, notochord cells, here representing the axial tissues.

In a recent 1-dimensional continuum model of elongation^13^, progenitor influx was assumed to be proportional to the cell density gradient at the pPSM-tailbud/progenitor domain (PD) border, which effectively ties elongation (which reduces cell density) with proportional compensation of new cells, allowing the elongation speed to be stable. Whether this progenitor influx regulation takes place *in vivo* remains to be tested, although it is known that other body axis tissues such as the elongating notochord can affect progenitor movement^2^. In this study, we examine how cell density of the pPSM regulates body axis elongation and correspondingly, its own cellular influx. This is achieved by a 3D mutli-tissue model that captures the mechanical coupling of the pPSM and axial tissues, and experimental tests that alter the elongation progress and pPSM cell density in the intact embryo. We further examine how FGF signalling intersects with this process, enabling cell density to serve as a dynamic regulator of the elongation speed.

## RESULTS

### 3D multi-tissue modeling suggests the balance between elongation and new cell influx underlies dynamic stability

To recapitulate 3D inter-tissue interactions that are important for elongation and cell density regulation, we developed agent (i.e., cell)-based models that follow the multi-tissue layout of the posterior body axis region. For the scaled 3D-model, we utilized PhysiCell, an open-source 3D modelling software^14,15^. The simulation construct (∼8k cells for the 3D model, comparable to the real tissue) contains a passive central axial tissue (representing the notochord and the neural tube) flanked by pPSM tissues, and a PD on the posterior midline (Figure 1B). This construct is further confined by rigid anterior, dorsal, ventral and lateral boundaries (representing somites, ectoderm, endoderm and lateral plate mesoderm, respectively, modelled as non-cellular), and a posterior boundary that is modelled as either passive movable non-PSM cells or free space. PhysiCell was chosen as it is a physics-based simulator, where cells are not constrained by lattice positions, and instead move independently between iterations, following the biomechanical forces they experience locally. This allows one to study emergent tissue-scale properties of multicellular dynamics in mesenchymal tissues such as the pPSM.

In the 3D model containing only the tissue geometrical layout and motile pPSM cells, we found that a confined pPSM undergoing expansion is sufficient to drive most of the tissue flow features observed in embryos, including convergence and elongation of both the PSM and the axial tissue (Figure 1B). A 2D equivalent of the 3D model produces similar results and allows addition of new cells to the PD^2^, therefore we used the much faster 2D model (∼300-800 cells) to scale up the number of independent simulations. With progenitor influx, we reproduced stable elongation and a sustained cell density in the pPSM (Figures 1C-D). Reducing this influx led to a fall of cell density and slowing of the elongation progress.

Using parameter sweeps, we also found that the cell activity of the pPSM (parameterized as cell-cell repulsion in the model) is the key factor for axis elongation under a constant initial cell density and in the absence of proliferation (Figure 1E). Thus, both the previous study^13^ and our results show that the space created by elongation (driven by pPSM cell activity) needs to be continuously filled with new active cells to sustain axis elongation, although the underlying mechanisms of progenitor influx regulation employed by both models are distinct and not mutually exclusive. The former depends on the progenitor movement following the cell density gradient at the pPSM/PD border which could become steeper as a result of pPSM expansion^13^. The latter works through inter-tissue mechanics where the expanding pPSM compresses the axial tissue to elongate and consequently push the midline PSM progenitor cells in the PD to move laterally into the pPSM^2,12^.

### Mechanical perturbations alter body axis length and pPSM cell density patterns

To alter cell density in the pPSM *in vivo* in a minimally-invasive manner, we extended the body axis of HH10-12 embryos by pulling and holding the tissue for 1-2 minutes from anterior to posterior (AP) at a contact point near the Area pellucida edge (Figure 2A). Upon release of the stress, the bulk of the body axis exhibits an elastic recoil to restore the tissue length. However, the posterior end retains ∼200µm of stable length increase, which is within the range of body axis length variation observed in unperturbed embryos at the same stages (Figure 2B). The posterior extraembryonic tissues exhibit even more prominent plastic deformation with a smaller recoil (Figure 2A). To exert the opposite effect on the axis length, we compressed embryos using tissue force microscopy (TiFM)^12^ for ∼5 min before retrieving the probe. Analogous to pulled embryos, this operation generated an initial recoil which stabilizes to a shortening of ∼100µm at the posterior end of the axis (Figure 2C). The posterior extraembryonic tissue was observed to creep anteriorly during compression and showed minimal recoil, consistent with its behaviour under pulling (Figure 2C). These results show that tissues along the AP axis have a graded transition from stiffer and elastic to softer and plastic materials, corresponding to the reported gradients of extracellular matrix expression and cell organization^1,16^.

**Figure 2.**
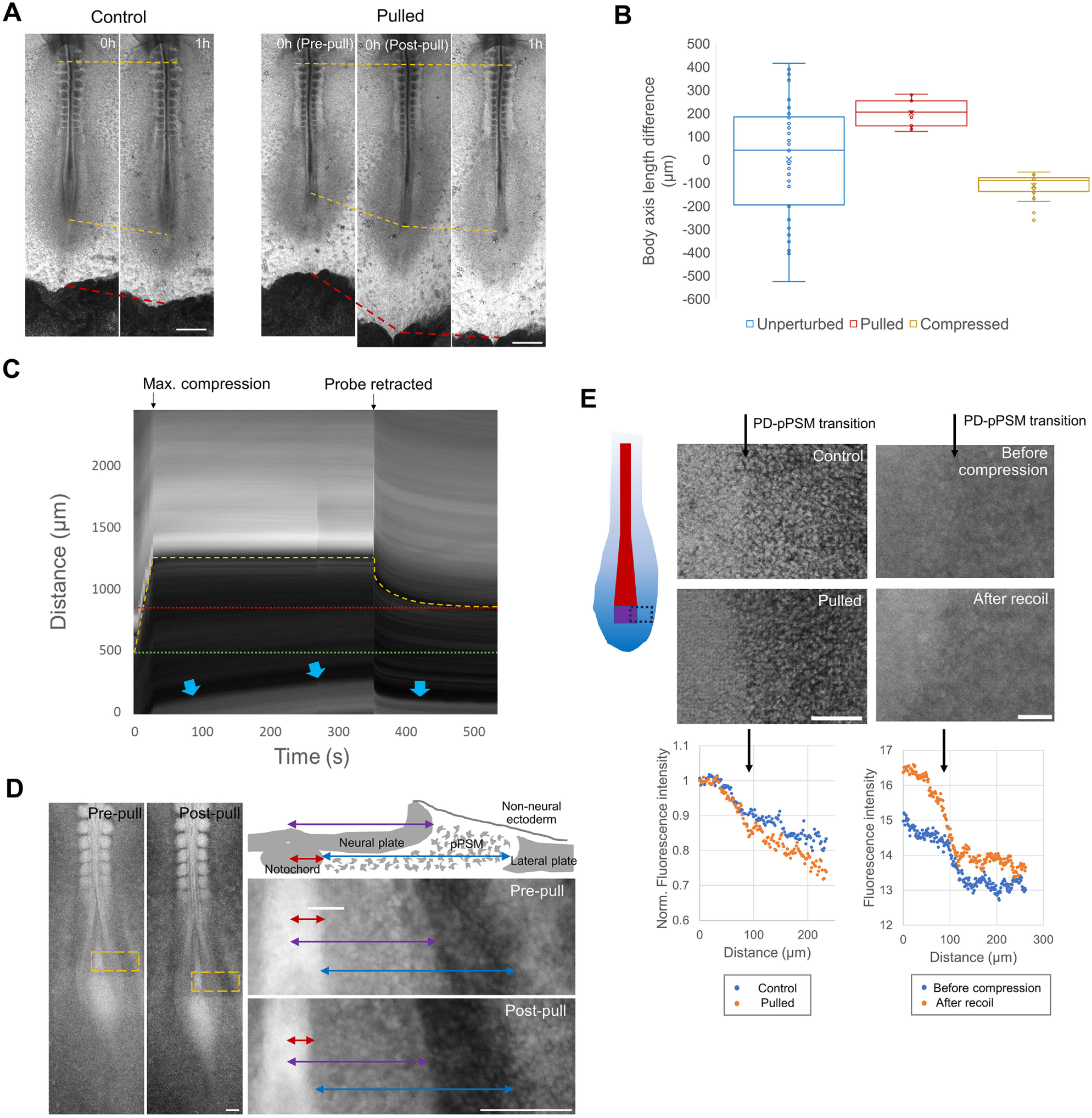
Tissue deformation in extended and shortened embryos. **A.** Deformation of the body axis and surrounding tissues after pulling. Ventral views with anterior to the top (same in other panels). The ends of the top yellow dashed lines mark a somite reference and the ends of the middle yellow dashed lines mark the end of the axis. The bottom red dashed lines mark the boundary of Area pellucida (light grey) and Area opaca (dark grey). Scale bars: 500µm. **B.** The variation of body axis lengths in unperturbed embryos (the difference from the population average, n=45) compared to the plastic length changes of pulling and compression experiments (the difference before and after, n=11,20, respectively). Shorter/compressed was defined as negative. **C.** Kymograph of a representative compressed embryo, anterior to the top. Dashed yellow line tracks the inserted foil and the slit-wound after its retraction. Red dotted line shows the stable wound location while green dotted line shows the initial position of the foil. Blue arrows highlight the Area pellucida and Area opaca boundary, which shows a creep behaviour during the holding stage and minimal recoil after probe retraction. **D.** GFP+ embryo pulled by a TiFM probe. Images show the GFP signal. Orange boxes on the embryo images (left) mark the regions of zoomed-in views (right). The schematic shows the cross-sectional arrangement of body axis tissues (right side). Red, purple and blue double arrows indicate the width of the notochord, neural plate and pPSM, respectively. Scale bars: 100µm. **E.** Density difference assessed by fluorescence intensity in GFP+ embryos at the PD-pPSM transition. The illustration shows the area imaged (dotted black box. red, notochord; purple, PD; graded blue, PSM, same for the following panels). Arrows mark the transition boundary on images and the plots. The pulling confocal images (maximum projection) use 2 different embryos, the fluorescent intensity was normalized to the PD before comparison. The compression images were taken with epifluorescence in the same embryo (2 min post probe retraction). Scale bars: 100µm.

To analyse tissue shape and cell density changes following pulling and compression, we imaged the Roslin Green Tg(CAGGs:eGFP)^17^ embryos that provide high contrasts at tissue boundaries through the fluorescence signal. We found that the pulled embryos show a longer, darker PSM with unchanged width while the neighbouring neural tube and notochord show a narrower tissue^12^ (Figure 2D). This suggests that extension of the mesenchymal pPSM leads to a decrease in cell density, whereas extension of the epithelial axial structures causes tissue narrowing, mimicking their respective normal, active modes (expansion of the pPSM and convergent extension of the axial tissues, respectively) of elongation^2,3,9,18^. We measured the medial-lateral cell density gradient across the PD-pPSM transition area using the fluorescence intensity and found that the gradient is steeper in pulled embryos (Figure 2E). In the compressed embryos, the cell density difference at the PD-pPSM boundary also becomes larger (Figure 2E).

To validate the cell density change in the pPSM after pulling, we carefully compared stage- and location-matched control and pulled embryos using confocal imaging (Figure 3A). By counting labelled nuclei in confocal slices of the mPSM and pPSM regions along the body axis, we confirmed a marked decrease of cell density in the extended pPSM, particularly near the posterior end/progenitor domain (PD) of the tissue (∼20% decrease, Figure 3B-D). Conversely, compressed embryos presented with a density increase in the pPSM (Figure 3C-D). From high-resolution confocal images of the pPSM of embryos fixed immediately after pulling and compression (Figure 3D), we observed that the density increase seen in compression is accounted for by a decrease in the gaps between cells and thus a decrease in extracellular space. On the other hand, in pulled embryos, the lower density is accompanied by an increase in extracellular space. In all cases, cells exhibit mesenchymal characteristics with irregular shapes and multiple protrusions, and randomly oriented nuclei. These results suggest that pPSM lengthens and shortens under the mechanical stress through changing the amount of extracellular fluid, leading to the condensing or dilution of cells.

**Figure 3.**
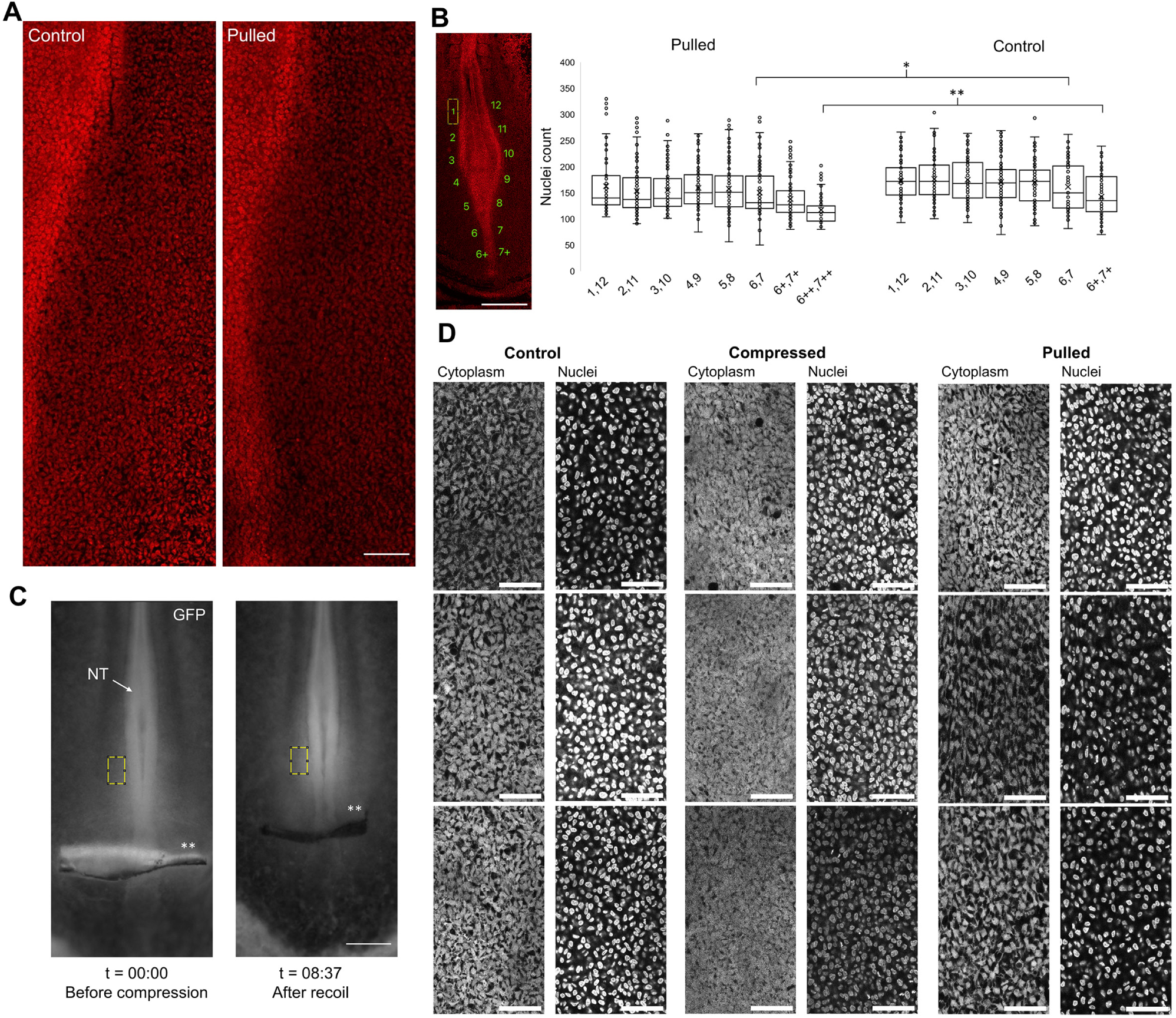
PSM cell density and extracellular space changes in extended and shortened embryos. **A.** Confocal images of the pPSM area on the right side of the axis. DAPI signal is shown in red. The denser regions to the left are the axial tissues and PD. Scale bar: 100µm. **B.** Cell counts of the areas marked in the image. Counting was done manually on 20 confocal slices through the PSM tissue per embryo (n=6 pulled embryos and n=4 control embryos) in equal areas of 100×200µm as shown with the numbered yellow box in the image. Additional pairs of posterior end areas (6+, 7+, 6++,7++) were counted to match the extended length in the pulled embryos with regions in control embryos. *p=0.037, **p=3e-7, 2 tailed t-tests. Scale bar: 500µm. **C.** Fluorescence change in compressed pPSM. Mean normalized (to undeformed extraembryonic tissue in the same image) fluorescence intensities in the tracked yellow boxes are 2.48 before compression and 3.74 after recoil, respectively. Asterisks mark the foil/wound location. Scale bar: 300µm. **D.** Representative confocal images of the pPSM of control, pulled and compressed embryos (3 of each). Cytoplasmic GFP signal shown on the left and Hoechst signal shown on the right. Dark space in the GFP images shows extracellular space. Scale bars: 50µm.

### Progenitor influx decreases with length increase and vice versa

To test how progenitor influx responds to cell density changes in the pPSM, we labelled the midline PSM progenitors in the PD by DiI injection and tracked their spread^2^ (Figure 4A). Surprisingly, the progenitor spread is reduced after pulling (Figure 4B) against the steeper density gradient (Figure 2E). This result shows that the feedback between elongation and cell influx does not occur at the PD-pPSM border as suggested previously^13^. Instead, our multi-tissue model suggests that the lower-density pPSM after pulling now produces a reduced stress (Figure 4C) for the axial tissues, which in turn push less strongly on the PD (Figure 4D), further reducing the progenitor influx into the pPSM (Figure 4E). It is unlikely that the reduced stress is due to changed FGF activity in pPSM cells as their motility (measured through the spreading of labelled pPSM cell clusters) remains unchanged in extended embryos (Figure 4B). This motility is known to be controlled by the FGF activity level in the pPSM and inhibiting it causes stalled elongation^1^. Conversely, progenitor influx was observed to increase in the shortened embryos (Figures 2A-B), following the steeper density gradient and further increasing cell density in the pPSM which had already been increased directly by the compression (Figure 2E). These results together show that cell density variation in the pPSM is not only uncompensated by the progenitor influx, but also further exacerbated, likely through the mechanical coupling of the pPSM and the axial tissues.

**Figure 4.**
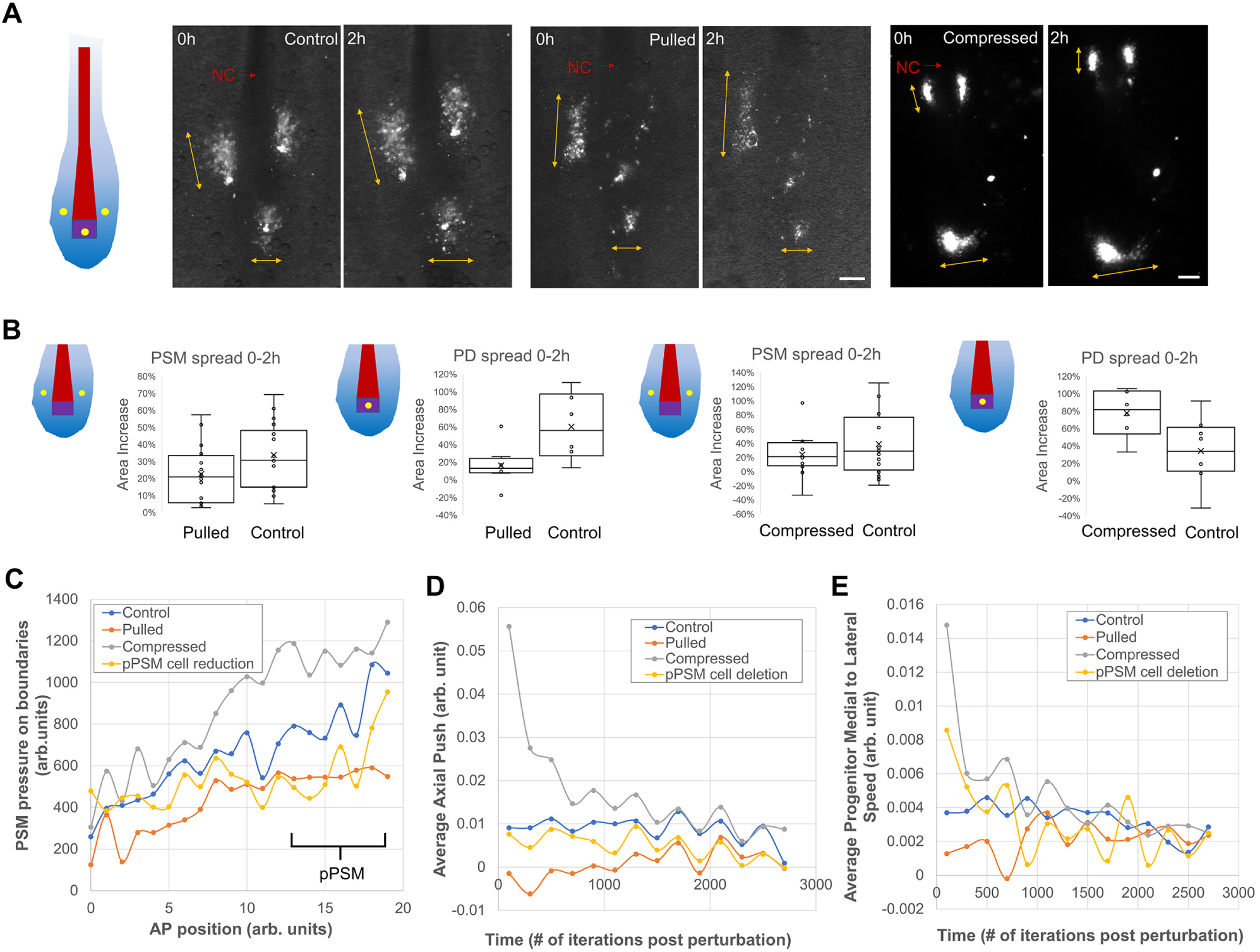
Progenitor cell influx in extended and shortened embryos. **A.** Tracking cell clusters by injected DiI in pPSM and PD (yellow spots in the illustration). Double arrows show the spread of the clusters. NC, notochord. Scale bars: 100µm. **B.** Cluster area (segmented around the edge for pulling experiments and calculated as a product of AP and LM spreads for compression experiments, respectively, due to different image qualities) changes over time. PSM spread pulled n=16, control n=18, p=0.076. PD spread pulled n=8, control n=6, p=0.019. PSM spread compressed n=12, control n=16, p=0.338. PD spread compressed n=6, control n=8, p=0.036. 2 tailed t-tests. **C.** Simulated pressure recorded along the AP axis by tallying the collisions and collision strengths of PSM cells with the boundaries of the cell field over time. Results were binned to one average value per unit length (the starting field length is 15 units, pPSM as defined here takes ∼5 posterior units up to the posterior-most axial cells). Curves show the average pressures of 10 simulations. Compression and pulling use a parameter of 80 (the same parameter was used in panels **D**,**E** and in the strong pull and compression conditions in Figures 5C-D). pPSM cell deletion causes pressure drop in the pPSM as expected. The pressure increase by compression also affects pPSM more prominently, while pulling causes a more global decrease. **D.** Simulated axial pushing force as measured by the A to P forces on the posterior-most axial cells. Results were binned to one average value per 200 iterations after introducing the perturbations. Curves show the average forces of 40 simulations. This force dynamics is directly associated with the elongation speed. A stronger but shorter-lasting increase in the compression is contrasted with a weaker but longer-lasting decrease in the pulling. **E.** Simulated PD cells were identified as PSM progenitors that reside on the midline posterior to the axial cells at any given iteration and their medial-lateral speeds were tallied and binned to one average value per 200 iterations after introducing the perturbations. Curves show the average cell flow of 40 simulations.

### Model simulations predict long-term length recovery via speed changes

To evaluate the consequences of the aggravated cell density change following pulling and compression, we simulated the effects of these perturbations by global (Figure 5A) and graded extension/compression from the posterior end (Figures 5B-D), and measured body axis elongation speeds over time. As expected from the significance of cell density in force generation, the compressed embryos with a higher density showed an immediate increase of elongation speed while the pulled ones showed stalling of elongation. Interestingly, this response corrects for the length changes introduced by the mechanical perturbations, restoring axis length to that of the control, unperturbed embryos. Furthermore, this correction completes within a timeframe such that the average elongation speed, taking into account the introduced length changes, is indistinguishable from controls. The model thus suggests that cell density dynamics in the system stabilize the elongation speed against fluctuations, and constrain the axis length in longer term.

**Figure 5.**
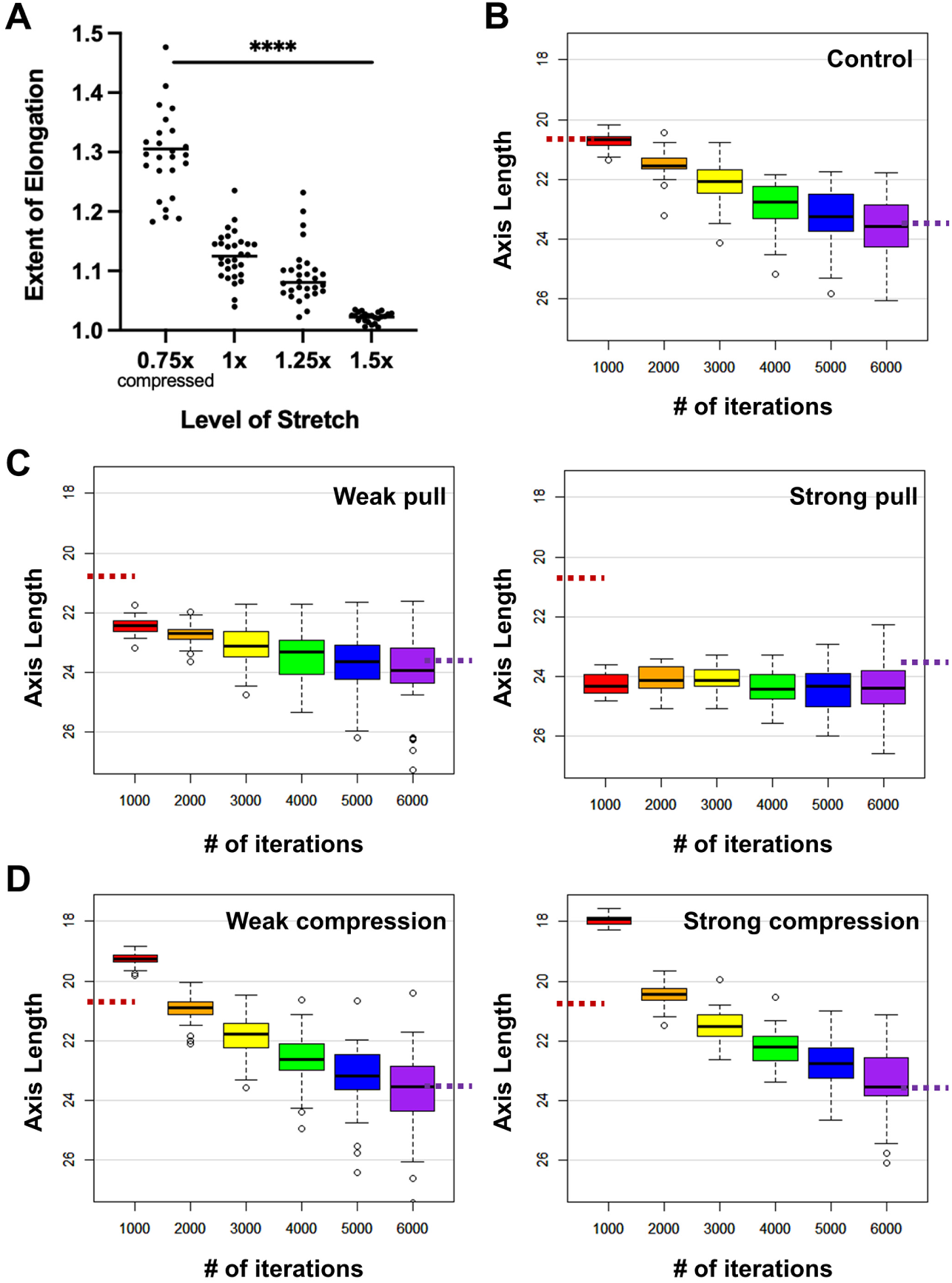
Simulated elongation responses to length perturbations. **A.** Extent of elongation (defined as the ratio between final simulated axial tissue length over the initial length immediately after the global stretch/compression, with no new cell additions during iterations). n=25-30 for each group. ****ordinary one-way ANOVA, F=55.98, p=1e-4. **B-D.** Elongation in the 2D model (including new cell additions) with local stretch/compression in the posterior end. n=40 simulations for each test. Dashed red and purple lines mark the average axis lengths in control embryos (panel **B**) at iteration 1000 and 6000, respectively.

### Cell density works synergistically with FGF activity to ensure the robustness of body axis length through dynamic speed regulation

To test this hypothesis experimentally, we tracked the elongation of pulled and compressed embryos. Strikingly, the pulled embryos showed recovery of body axis length by slowing down elongation immediately after pulling (Figures 6A, C). This decrease is seen both when comparing to control embryos (Figure 6C) and to the same embryos in the 2.5h before pulling (Figure 6E). The speed difference between controls and perturbed embryos corrects for the initial difference in length over time. Conversely, in compressed embryos, elongation speeds increase significantly in the first hour restoring the length to normal, despite the open wound in the posterior created by the insertion of TiFM probes (Figures 6B, D). In both cases, although the speed of body axis elongation changes, segmentation proceeds at the normal rate (Figures 6F), suggesting that pulling and compression do not affect the segmentation clock. These results show that body axis length and average elongation speed are robust properties of the system, possibly relying on the negative feedback through cell density and inter-tissue mechanics to react quickly to length deviations.

**Figure 6.**
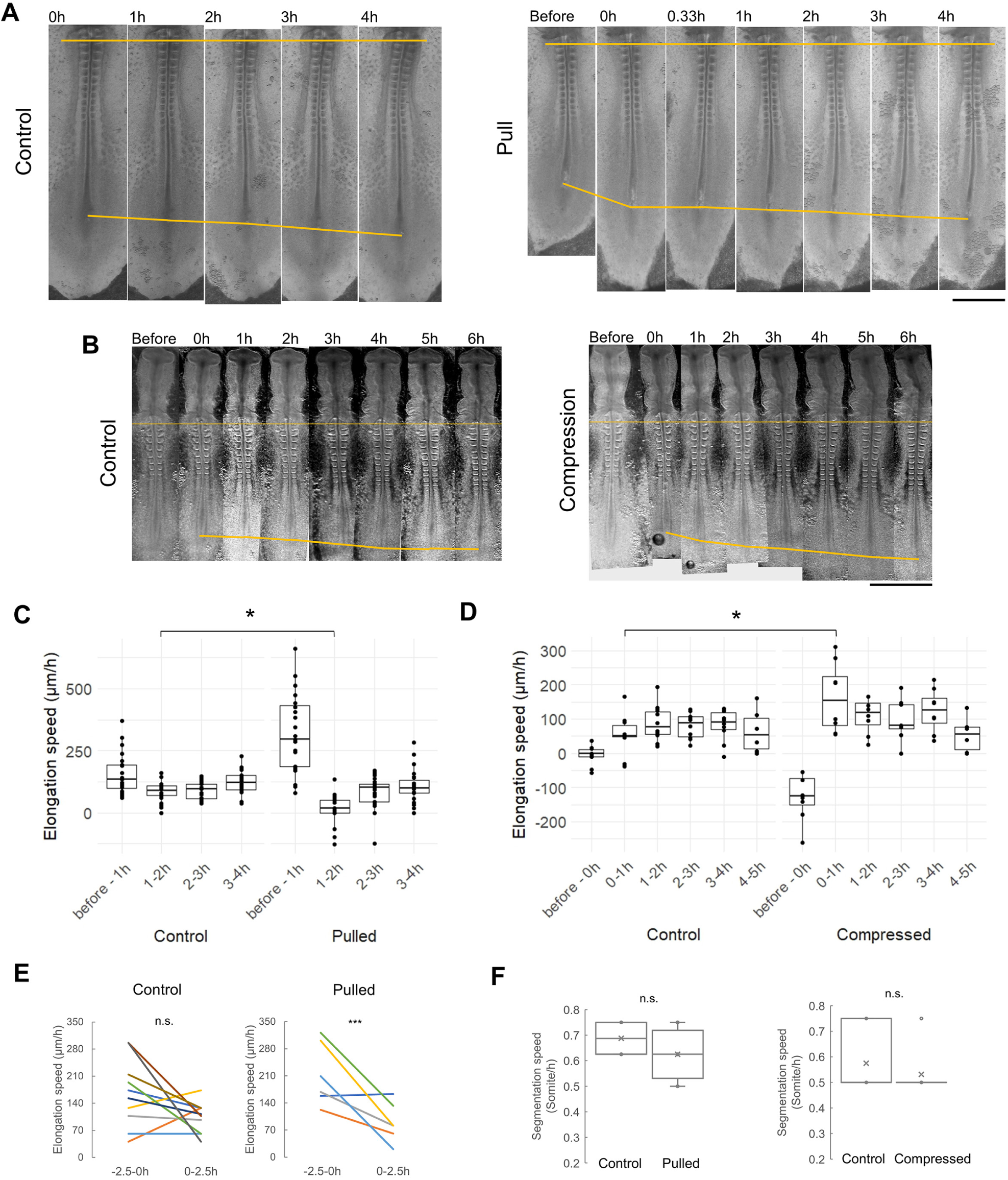
Elongation speed response and robustness to length changes. A-B. Representative embryo changes after pulling (**A**) and compression (**B**). The ends of the top yellow lines mark a somite reference and the ends of the bottom yellow lines mark the end of the axis. Scale bars: 1mm. **C-D.** Elongation speed dynamics following pulling (**C**, control n=24, pulled n=23, *p= 2e-5, 2 tailed t-tests) and compression (**D**, control n=10, compressed n=8, for 4-5h n=6 for control and compressed, *p=0.011, 2 tailed t-tests). **E.** Average elongation speeds in the 2.5h prior to pulling and 2.5h after. n.s., p=0.092, ***p=0.019, paired 2-tailed t-tests. **F.** Segmentation speed. n=8 for controls and n=12 for the pulled (n.s., p=0.118, 2 tailed t-tests).

As the acceleration and deceleration of elongation in response to cell density change should require the activity of pPSM cells under FGF signalling, we tested the relationship between FGF signalling and density changes under our perturbations. We first imaged fixed embryos after treatment with an FGF signalling inhibitor (PD173074) stained for phospho-MAPK (pMAPK, a downstream indicator of FGF signalling^10^). We measured the average intensity of the pMAPK signal along the PSM using a “U”-shaped mask (Figure 7A). The pMAPK signal was variable among individual samples but on average a posterior to anterior gradient can be observed, which diminishes under PD173074 treatment (Figure 7B), consistent with previous studies^1,10^. To evaluate how pulling and compressing affect this gradient, we measured pMAPK levels immediately after the perturbations. In compressed embryos, the gradient observed in controls is maintained (Figure 7C) and appears steeper on average. In pulled embryos, the gradient appears flattened on a higher level (Figure 7C). These results are consistent with the idea that the profile of FGF signalling gradient simply follows with pPSM length changes and is not directly altered by the mechanical perturbations, which are also consistent with the unchanged cell spread/motility observed in both extended and shortened pPSMs (Figure 4B).

**Figure 7.**
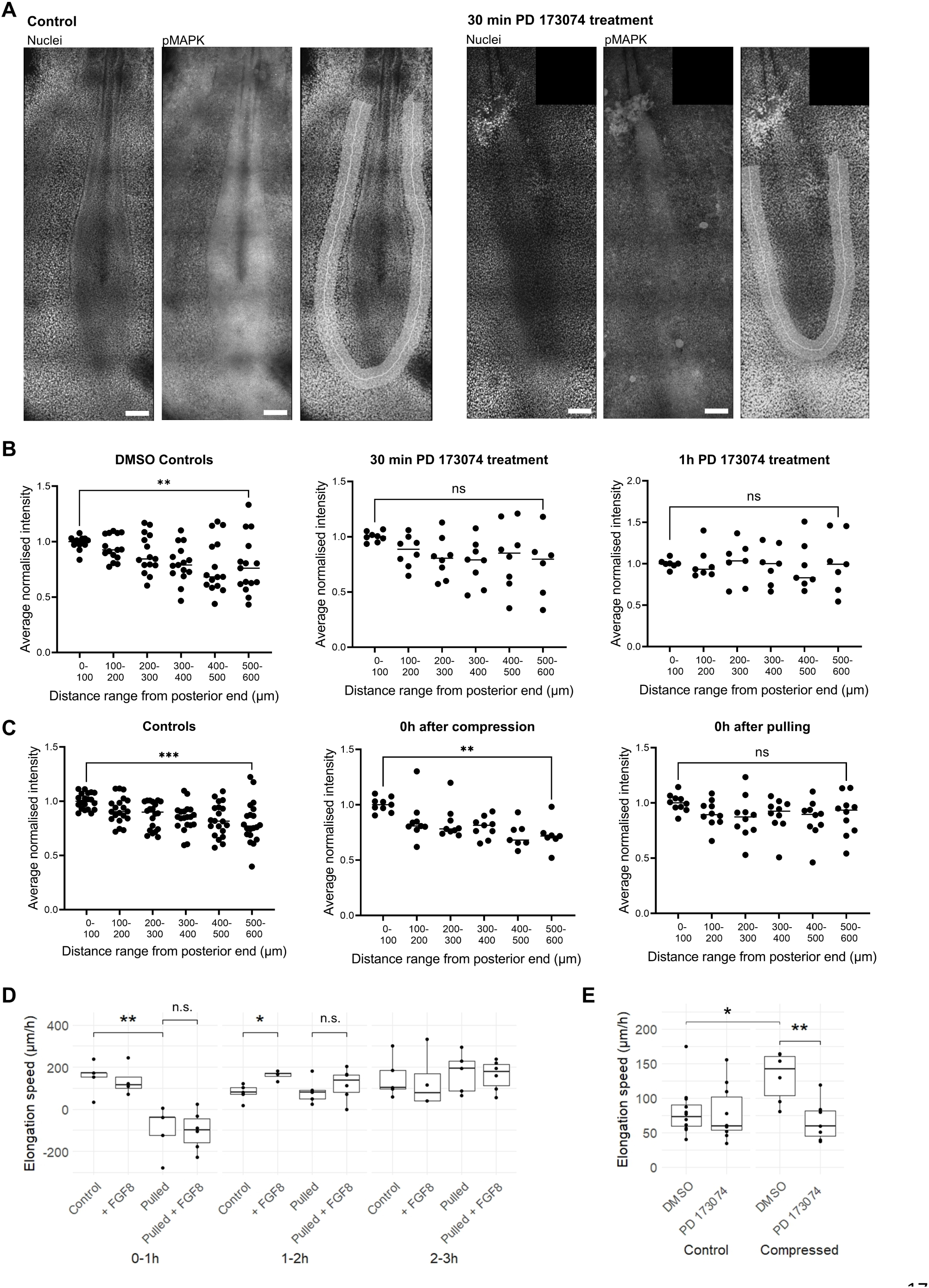
Integration of FGF signalling and elongation speed response. **A.** Representative confocal images of control embryos and embryos treated with the FGF inhibitor PD173074 for 30 min. Hoechst signal shown on the left, pMAPK signal shown on the middle, Hoechst signal with PSM shape drawn for intensity analysis (using Fiji) shown on the right. Regularly appearing, blurry dark horizontal lines are artefacts of the stitching of tiled confocal image edges. Scale bar: 100μm. **B.** Average fluorescence intensity (normalised to the average intensity of the most posterior point +100μm on either side, same in panel **C**) of pMAPK for controls (DMSO) and treated with PD173074 for 30 min and 1h. n=15 (8 embryos) for DMSO Controls, n=8 (4 embryos) for 30 min PD173074, n=7 (4 embryos) for 1h PD173074 (**p=0.0089, 30 min: n.s. p=0.058, 1h: n.s. p=0.95, paired t-test applied to samples for which there are datapoints at the two ends as not all pPSM images reach 600μm, same in panel **C**, n=6 for 30 min PD173074). **C.** Average fluorescence intensity of pMAPK for controls, after pulling and after compression. n=20 (10 embryos) for controls, n=9 for compressed (5 embryos), n=10 for pulled (5 embryos) (***p=3e-4, **p= 0.0017, n.s. p=0.11, paired t-tests, n=7 for compressed). **D.** Elongation speed dynamics following pulling + FGF8 addition. n=5 for control and pulled, n=4 for +FGF8 and n=6 for pulled + FGF8 (0-1h: **p=3e-3, n.s. p=0.913; 1-2h: *p=7e-3, n.s. p=0.431; 2 tailed t-tests). **E.** Average elongation speed over 4 hours after compression + FGF inhibition. n=12 for control (DMSO), n=10 for FGF inhibitor (PD173064), n=6 for compressed DMSO and n=7 for PD173074 (*p=0.011, **p=4e-3, 2 tailed t-tests).

To further analyse the relationship between cell density and FGF signalling in elongation speed regulation, we next measured elongation speeds in the hours following pulling while adding FGF protein to the posterior body. Unlike the control embryos that showed a transient boost of speed during 1-2 hours post treatment, the pulled embryos did not show a speed increase (Figure 7D), consistent with the idea that a certain cell density is required for active cells to generate the stresses for elongation. Conversely, the acceleration of compressed embryos was abolished in the presence of PD173074 (Figure 7E), indicating the shortened axis requires cell activities to produce the corrective response despite having a higher density pPSM. These results show that FGF activity and cell density work cooperatively to regulate elongation speed during body axis formation, providing a driver and a constrainer, respectively (Figure 1A).

## DISCUSSION

How pPSM cells combine density and FGF activity to generate the tissue level expansive force warrants further investigation. As a mesenchymal tissue, pPSM is found to contain cells that are protrusive and mobile, who exchange neighbors regularly and do not form stable junctions with others^1^. A hypothesis considering this cell state is that the motility of the cells creates diffusion-like behaviour that expands the pPSM^1,2,13^. Cell density in this case will be directly indicative of the expansion potential, as also modelled in this study. The physical basis of this analogy, however, is unclear as cells move in an over-dampened environment and their movements do not translate directly to pressures on each other or the tissue boundaries. A more recent hypothesis suggests the ECM, specifically hyaluronic acid, draws in water to swell the pPSM^3^. Cell density in this case will be proportional to hyaluronic acid concentration (assuming cells produce it at similar rates) thereby tissue expansion. Our pulling and compression experiments of the pPSM support this possibility as we do not observe major changes in cells (shape, orientation, motility, or FGF activity), instead we observe the tissue volume changes to be mainly in the extracellular space, suggesting water gain/loss following shortening/extension. Indeed, an optimal hydration level through this mechanism may link cellular regulation to tissue forces to define the elongation speed. This will be important to test in future combining the mechanical approaches here with osmotic control and fine measurements of tissue hydration. It will also be necessary to examine more broadly the genetic and biochemical changes to the cells and tissue following cell density changes, and to develop tools that can alter cell density non-invasively without changing the pPSM tissue shape at the same time.

Looking beyond the pPSM, it is important to note that body axis elongation is a multi-tissue process. While the tissue layout and inter-tissue normal forces between the pPSM and the axial tissues have been modelled here and in our previous study^2^, other mechanical factors such as the shear stress between tissues and the rheological patterns^3,16^ of the pPSM remain to be investigated. One notable observation is that when we perform pulling, a narrow probe anchored on the axis end was sufficient to extend not only the neural tube but also the PSM and lateral plate tissues. In contrast, when we compress with a narrow probe only the axial tissues will shorten while the pPSM and other tissues do not follow.

This required us to use a much wider probe that can also compress pPSM directly to achieve pPSM shortening, suggesting that tissues are coupled by shear stress in one direction (A to P) but not the other (P to A). Combining TiFM-based stress measurement and molecular perturbations of inter-tissue connections such as collagen^19^ and fibronectin^20^, could lead to new insights in tissue mechanical coupling during elongation.

More generally, the mechanisms of morphogenetic speed control remain poorly understood. Our work reveals a simple strategy for expanding mesenchymal tissues. Progress is driven - but also limited - by cell density, where the speed is constrained by the timescale that arises from the interactions between cell density regulators. This mechanism provides stability and long-term robustness that are also tunable by active factors (such as signalling). In the context of vertebrate body axis formation, this mechanism, in conjunction with the segmentation clock, ensures sequential somite formation at consistent sizes and appropriate number, with an invariant axis length for a reproducible body plan.

## LIMITATIONS OF THE STUDY

Finer dissection of the relative contribution of cell density and FGF activity to the elongation speed is not possible due to the normal variability associated with body axis lengths and elongation speeds found in the chicken embryos. In the TiFM experiments which are low-throughput, embryos experience a period of low temperatures which likely caused a lower average elongation speed in control embryos, making the effect of further perturbations (e.g., PD173074) difficult to detect. Similarly, in FGF inhibited embryos that will have low baseline speeds, the effect of compression is challenging to measure, preventing a test of whether compression-induced acceleration is additive to or permitted by the processes under FGF signalling.

## ACKNOWLEDGEMENT

The authors thank C.U. Chan, O. Pourquie, N.J. Lawrence, S. Kato and members of the Xiong lab for reagents, technical assistance, and comments; R. Heiland for training and debugging with the PhysiCell Studio software. This study is supported by a Wellcome Trust / Royal Society Sir Henry Dale Fellowship (215439/Z/19/Z) and an UKRI-EPSRC Frontier Research Grant (EP/X023761/1, originally selected as an ERC Starting Grant) to F.X. C.L. acknowledges a China Scholarship Council (CSC) scholarship (201906240004). A.M. acknowledges a Cambridge Centre for Physical Biology (CPB) summer scholarship.

## AUTHOR CONTRIBUTIONS

C.L., J.M.N.V., C.C.J.J. and F.X. designed the project. C.L. and J.M.N.V. performed the embryo experiments, A.M. and F.X. contributed to the embryo experiments. C.C.J.J., A.M. and F.X. performed the simulations. J.M.N.V. and F.X. prepared the manuscript. All authors analysed the data and contributed to the manuscript.

**The authors declare no competing interests.**

## STAR METHODS

### RESOURCE AVAILABILITY

#### Lead Contact

Further information and requests for resources and reagents should be directed to and will be fulfilled by the Lead Contact, Fengzhu Xiong.

#### Materials Availability

This study did not generate new unique reagents.

#### Data and Code Availability

The published article includes most datasets generated during this study. Additional replicas and source datasets are available upon request. The computational model is publicly accessible via the link provided in the Key Resource Table.

### EXPERIMENTAL MODEL AND SUBJECT DETAILS

#### Chicken embryos

Wild type fertilized chicken (*gallus gallus*) eggs were supplied by MedEgg Inc. Transgenic eggs were supplied by the National Avian Research Facility (NARF) at University of Edinburgh. Eggs are stored in a 14°C fridge and incubated under 37.5°C ∼45% humidity incubators (Brinsea). No animal protocol is required for the embryonic stages studied under the UK Animals (Scientific Procedures) Act 1986 (under 2 weeks, or 2/3 of gestation time for chicken).

### METHOD DETAILS

#### Embryo preparation

Embryos were staged following the Hamburger and Hamilton (HH) table^21^ following extraction from the egg using the Early Chick (EC) culture protocol^22^. The embryos were maintained in a slide box with wet paper towels in incubators at all times except when snapshot images (<2min per embryo) were taken and when injection, pulling and compression experiments were performed either manually or on the TiFM system under room temperature. DiI injections were performed with a sharp-tipped glass needle with a Nanoject microinjector or mouth pipetting at 0.5mg/ml (2.5mg/ml DiI in ethanol diluted in Ringer’s solution or PBS immediately prior to needle loading). The needle enters from the ventral side targeting the PD and the pPSM. The initial injected spots were allowed to diffuse for 2-4 hours to allow trackable single cells to appear around the spot, before pulling and compression perturbations were performed to the tissue. FGF signalling was inhibited by treating the embryos with 50 µl of a 4 µM solution of the FGF inhibitor PD173074 (Bio-techne). To enhance FGF signalling, 50 µl solution of FGF8 recombinant mouse protein (424-FC-025/CF, R&D systems) at a concentration of 1 µg/ml was dropped by a pipette on the ventral surface of the embryo.

#### Manual and TiFM pulling and compression

HH10-12 embryos prepared on the EC culture plates were used. The embryos were under incubation prior to the experiments (conducted at room temperature) and returned to incubator after the recoil stopped (∼2min, embryos left in room temperature for extended times do not show further recoil or elongation). Pulling experiments were performed both manually as well as on the TiFM. Compression experiments were performed on the TiFM alone. For manual pulling, a tungsten rod (∼50µm diameter) was installed on a surgical needle holder (Fine Science Tools) and brought to contact with the embryo at the boundary between Area opaca and Area pellucida at the axial midline point. Gently, without creating tearing damage, the tissue was pulled posteriorly with visible strain of the body axis. A ∼30% strain from head to tail was achieved gradually and maintained for 1-2 minutes before retracting the rod. The tissue showed a recoil that stopped after 1 minute, retaining ∼100-200µm of posterior elongation (∼4% strain). For TiFM pulling, a cantilever probe was inserted into the posterior cells to perform the loading, mimicking the manual protocol to increase consistency. For compression experiments, as individual tissues tend to decouple when the compressed interface between the probe and the tissues is small, a tailored piece of aluminium foil that covers both pPSM cross-sections and the axial tissues was used. The foil was inserted just posterior to the PSM progenitor domain (PD). The axis was held for 5-6 minutes after ∼600µm of posterior to anterior foil movement and then the foil was directly retracted from the embryo. A thin slit wound was left in the posterior cells which normally healed later. The recoils of the tissue were recorded in timelapse movies and the movement was fit with a stress relaxation model to confirm the presence of a long-term plastic length change (only embryos that showed at least 50µm (∼1.5% strain) of shortening were further analysed). The embryos were then returned to the incubator and imaged regularly to measure axis elongation and cell spreading if labelled.

#### Immunostaining

Embryos were fixed by incubation with 4% paraformaldehyde overnight at 4°C and permeabilised by incubation with PBS with Triton X-100 (PBST; 0.5% Triton X-100) for 30 min at RT. Following 1h incubation at RT in blocking solution containing 4% Donkey Serum (D9663, Sigma-Aldrich) in PBS, embryos were incubated for 1-2 days with primary antibody, anti-dpERK (9101, Cell Signaling Technology), diluted 1:50 in blocking solution at 4°C. The primary antibody was washed off with PBS and the embryos were incubated with secondary antibody, donkey anti-rabbit IgG H&L (Alexa Fluor 594) (ab150076, Abcam) or donkey anti-rabbit IgG H&L (Alexa Fluor 647) (ab150075, Abcam), diluted 1:400 in blocking solution overnight. Hoechst (H3570, Invitrogen) (1:1000 dilution) was added on the final day for 10 min at RT. Upon nuclear staining with Hoechst, embryos were washed in PBS and mounted using ProLong Glass Antifade Mountant (P36982, Invitrogen) according to the manufacturer’s instructions.

#### Imaging

Snapshots of the embryos were taken with a stereomicroscope (MZ10F, Leica). Confocal images were taken with a laser scanning microscope with a 40X objective (on a Leica SP8). For live imaging on the TiFM, cultured embryos were transferred to a 35mm glass bottom dish (VWR) with a thin layer of albumen (200µl) and imaged using a Zeiss Axio Observer 7 microscope base as part of the TiFM system with a 5x or 10x objective at room temperature.

#### Data analysis

Images and movies are processed by ImageJ (NIH) and Powerpoint (Microsoft). Scale bars are first set with control images with objects of known sizes. To measure cell density, high resolution confocal Z-stacks of DAPI staining were used for manual nuclei count in ImageJ. For transgenic fluorescent embryos, the fluorescent signal from the PSM and PD tissue was used as an approximate measure of cell density, which was validated by DAPI counts. To measure cell spreading, DiI-injected tissue areas in the pPSM and the PD were marked as ROIs at different points with trackable cells forming the edges. The areas of these ROIs were then compared to yield a percentage change over time. To measure the elongation speed, the distance between a fixed somite pair (somites 2-5) and the posterior end of the body axis was taken. DiI was injected to mark the posterior end to minimize measurement inaccuracies due to tissue deformation after pulling. To measure fluorescence intensity of the pMAPK signal, a mask (ROI) for the “U”-shaped PSM was drawn on the maximum projection image and the intensities along the posterior to anterior axis under the mask were collected. Results were plotted with statistical tests performed using Excel (Microsoft) and custom R code or GraphPad Prism 9.4.1 (GraphPad Software, www.graphpad.com).

#### Modeling

Models were constructed and analyzed in Matlab (Mathworks), PhysiCell (including the interactive PhysiCell Studio). Detailed design and rationales are provided below. The models can be downloaded following the link in the Key Resource Table.

PhysiCell is an open-source, agent-based 3D cell simulator for mechanistic modeling of multicellular systems as previously described^15^. In this work, the mechanics module of the simulator was used to implement four main mechanical interaction parameters: cell-cell adhesion, cell-cell repulsion, cell-boundary adhesion and cell-boundary repulsion. The model resolves the net force resulted from these interactions and updates cell movement per iteration. The mechanical equation shown below relates the cell’s current velocity *v_i_* as a function of these terms and a movement term *v_i,mot_*, which describes active cell motility. Cell positions are *x_i_*. *ϕ_n,R(x)_* is an adhesion interaction potential, *ψ_n,R(x)_* is a repulsion interaction potential function. *c^i^_cca_* and *c^i^_ccr_* are the ith cell’s cell-cell adhesion and repulsion parameters, *R_i_* is its radius, and *R_i,A_* is its maximum adhesion distance. *d(x)* is the cell’s distance to the nearest boundary and *n(x)* is a unit vector normal to the boundary.

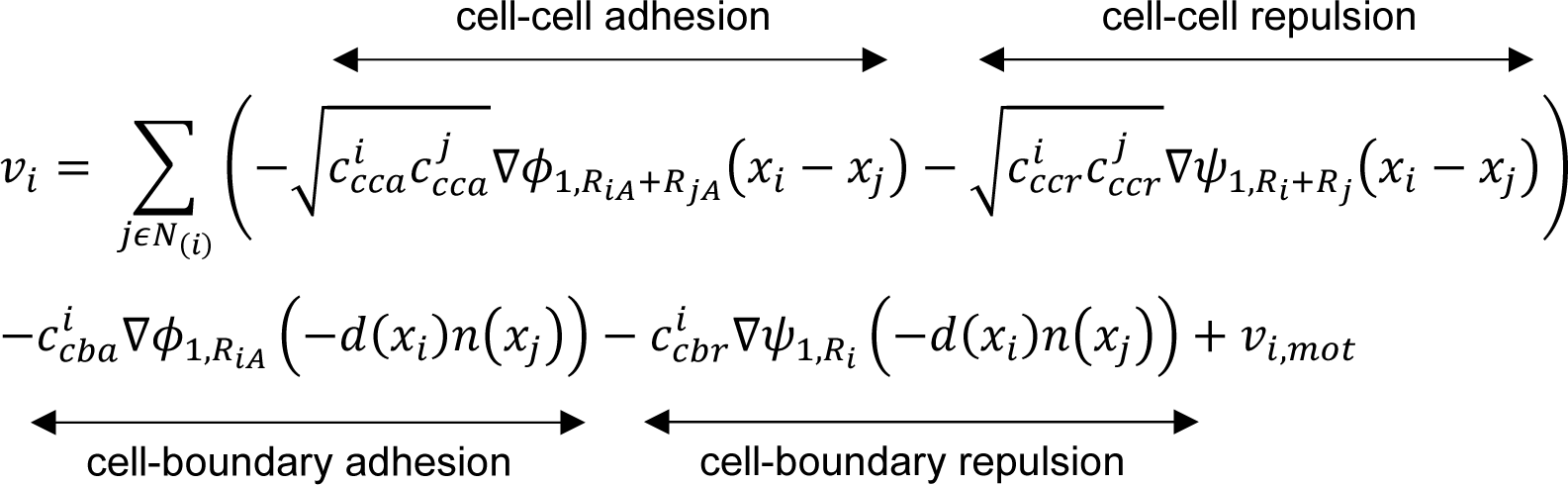

Simulations were run for a total of 7200 virtual minutes on a field of 7920 cells, using the following parameters:

| Cell type | Motility<br>(directionless) | Relative<br>Adhesion | Relative<br>Repulsion | Adhesion<br>Distance |
| --- | --- | --- | --- | --- |
| Axial | 0 | 2.5 | 10 | 1.1 |
| Presomitic Mesoderm | 1.2 | 0.05 | 10 | 1.25 |
| Posterior Cells | 0 | 1 | 10 | 1.1 |

For simplicity, axial cells including the notochord and neural tube were considered as one passive tissue (the motility of these cells was 0) that only deforms by the forces from the flanking PSM. The posterior cells were also simulated as non-migratory and passive which were displaced by an intermediate repulsion from the axial cells and the PSM cells. It is worth noting that these assumptions ignore the active intercalations of the axial cells as the tissues elongate^2^ and also the medial to lateral movements of the posterior cells. No assumptions about cell cycle, cell apoptosis, or tissue microenvironment was included in the model.

The output of the 3D model was compared with a simpler 2D model previously described^2^. No significant qualitative differences of tissue and cell behaviors were observed. The 2D model was then used to perform computationally demanding parameter explorations to understand how the various mechanical parameters affect axis elongation and convergence. The parameter space explored are as follows:

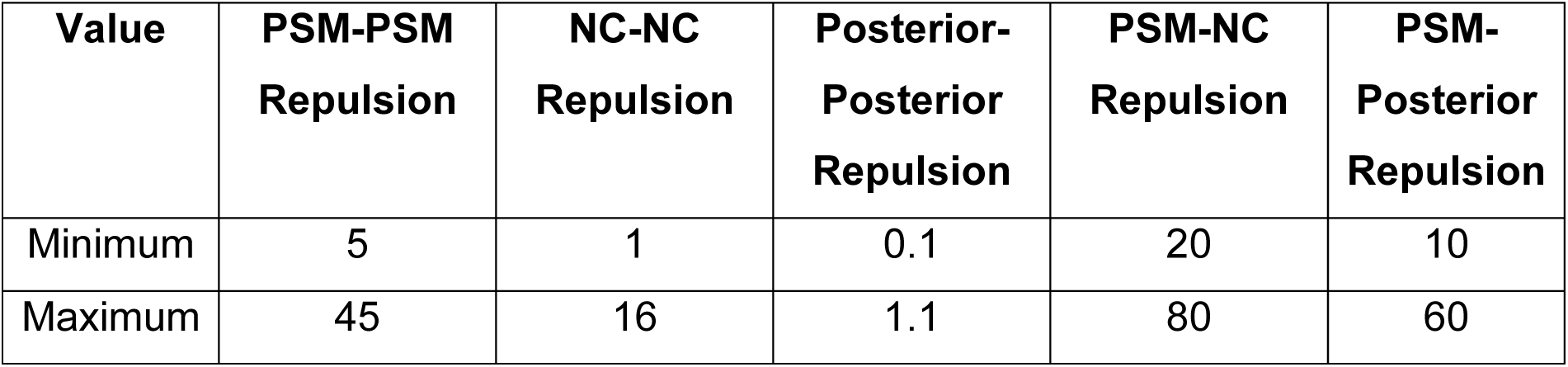

| **Value** | **PSM-PSM**<br>**Repulsion** | **NC-NC**<br>**Repulsion** | **Posterior-**<br>**Posterior**<br>**Repulsion** | **PSM-NC**<br>**Repulsion** | **PSM-**<br>**Posterior**<br>**Repulsion** |
| --- | --- | --- | --- | --- | --- |
| Minimum | 5 | 1 | 0.1 | 20 | 10 |
| Maximum | 45 | 16 | 1.1 | 80 | 60 |

To assist the parameter space coverage, an unbiased, unsupervised shallow neural network (created using Matlab, Mathworks) that did not require prior knowledge about the output targets was utilized to handle a total of 9193 outputs from the simplified 2D model. The 5 variables, PSM-PSM repulsion, NC-NC repulsion, Posterior-Posterior repulsion, PSM-NC repulsion and PSM-Posterior repulsion were used as inputs (table below), and the length and width of the axial tissue (determined from a minimum bounding box containing NC cells) were used as targets for the model. Both inputs and outputs were scaled before being used for training. 70% of the samples were used as training data, 15% as validation, and 15% as testing.

To model progenitor influx and axis length changes, we used a modified version of the previously described 2D model^2^ in Matlab (Mathworks). To simulate pulling and compression, the cell positions were adjusted as compared to control in either global deformation or a data-matched decaying deformation from posterior to anterior. In the former, the whole simulated field of cells was compacted or stretched equally along the AP axis. In the latter, stretching and compression were implemented in an exponential manner. The posterior cells were displaced more which was consistent with the experimental data probably because the tissues closer to the pulling rod or compressing foil were less elastic. The posterior 75% of the cells in the simulation field were pulled or compressed to reach a ±10%-15% total strain, to be consistent with the experimental data. Body axis elongation (measured by the average position of the 4 posterior-most axial cells) was then followed in the simulation for 6000 iterations per independent simulation. Groups of 10-40 simulations per control or test group were performed.

## QUANTIFICATION AND STATISTICAL ANALYSIS

### Statistical analysis summary

Quantitative results were categorized into experimental groups and Student’s t-test is performed with Excel (Microsoft), custom R code and GraphPad Prism 9.4.1 (GraphPad Software, www.graphpad.com) to compare the results with p<0.05 considered significant difference. The test type, value of n, definition of center and variation are all defined in each figure legend in the manuscript.

### Statistical considerations in study design

Sample size: For quantitative comparisons (e.g., elongation speed), at least 5 samples per group were used per experiment. This sample size is in line with the expectations in the field and is optimal for completing multiple groups of experiments in one sitting, taking into consideration the time required for embryo preparation and surgical operations.

Replications were performed to confirm the results. Before applying t-tests, the measurements were normalized to the mean and pooled for a chi square goodness of fit test. The tests suggest that the variabilities observed in axis lengths, elongation speeds, cell spread, nuclei count and fluorescence intensities do not come from a distribution significantly different (p>0.05) from a normal distribution. Data exclusions: one compressed sample was excluded from elongation analysis as its segmentation rate was low and thus it was deemed not to be developing properly; one sample was excluded from the pMAPK intensity analysis due to the region of interest being shadowed likely as a result of debris during mounting obscuring the signal. In sample preparation, embryos that showed unspecific developmental abnormalities were discarded. Replication: All wet-lab experiments reported have been replicated at least once. Randomization and Blinding: After an initial health screen, the investigators sort embryos into control and experimental groups randomly. Blinding is not possible for pulling and compression experiments as the perturbations leave clear differences of tissue morphology.

KEY RESOURCES TABLE

KEY RESOURCES TABLE
| REAGENT or RESOURCE | SOURCE | IDENTIFIER |
| --- | --- | --- |
| Chemicals, Peptides, and Recombinant Proteins |  |  |
| BactoAgar | Sigma-Aldrich | A5306 |
| PD173074 | Biotechne | 3044 |
| Dil | Sigma-Aldrich | 42364 |
| FGF8 recombinant mouse protein | R&D systems | 424-FC-025/CF |
| anti-dpERK | Cell Signaling Technology | 9101 |
| Experimental Models: Organisms/Strains |  |  |
| Wild Type Chicken Eggs | MedEggs Inc. | n/a |
| Tg(CAG-GFP) Chicken Eggs | University of Edinburgh; (McGrew et al., 2008) | n/a |
| Software and Algorithms |  |  |
| Fiji | NIH | https://fiji.sc/ |
| Matlab (R2022a) | Mathworks | https://www.mathworks.com/products/matlab.html |
| PhysiCell and PhysiCell Studio | Indiana University, Bloomington; (Ghaffarizadeh et al., 2018; Heiland et al., 2023) | http://physicell.org/<br>https://github.com/PhysiCell-Tools/PhysiCell-Studio |
| Agent-based mutli-tissue elongation model (in PhysiCell and Matlab) | this paper | https://github.com/ChrisChanJ/PhysiCell2024<br>https://github.com/xionglab/project_elongation |

